# Olfactory bulb and cortex activity reflects subjective odor intensity perception rather than concentration

**DOI:** 10.1101/2025.05.26.656087

**Authors:** Frans Nordén, Irene Zanettin, Mikael Lundqvist, Artin Arshamian, Johan N. Lundström

## Abstract

Understanding stimulus intensity processing is fundamental in sensory science, yet this question remains largely unexplored in human olfaction. We investigated how the human olfactory bulb (OB) and piriform cortex (PC) process odor concentration versus subjective perceived intensity. We demonstrate that OB-PC network oscillatory dynamics are predominantly driven by perceived intensity, not physical concentration. The OB initially processes and communicates perceived intensity to the PC via early gamma-band oscillations (bottom-up feedback). The PC then refines and sends this percept back to the OB via later beta-band oscillations (top-down feedback), updating the OB’s gamma activity for subsequent odorants. Critically, analyses of phase-amplitude coupling and beta burst activity demonstrate that transient beta patterns from the PC update OB gamma activity, providing the OB with an updated internal representation of the odor percept. These results reveal an oscillatory mechanism by which the olfactory system maintains perceptual constancy and adaptability despite fluctuations in environmental odor concentrations.

## Introduction

A core function of all sensory systems is to decide a stimulus intensity. This basic perceptual processing, where stimulus volume is transformed into perceived intensity, enables us to differentiate a touch from a punch, a whisper from a scream, a candle from the sun, and a whiff from a stench. Today, we have a detailed mechanistic understanding of how the human auditory and visual systems encode both physical stimulus properties (e.g., volume and luminance) and their perceived perceptual counterparts (e.g., loudness and brightness). This is not the case for the human olfactory system where nearly all our knowledge of how the brain process odor intensity comes from animal studies, which primarily focuses on stimulus concentration because directly asking animal models for subjective trial-by-trial ratings of perceived odor intensity is not feasible.

The lack of trial-specific information presents a challenge given that perceived odor intensity is not a direct reflection of the physical concentration of odorant molecules. Rather, it represents a sophisticated perceptual construct in which the brain’s early olfactory structures transform the information from the raw chemical input, received from odor receptors, into subjective experience of odor intensity^1^. While stimulus concentration serves as the initial sensory signal, perceived intensity exhibits an imperfect logarithmic relationship to concentration^1^. This perceptual transformation is mediated by dynamic neural processes emerging from the interplay between bottom-up sensory input and top-down modulation^2^.

Studies in non-human mammalian models assessing how the brain process odor intensity has, to date, primarily focused on spike pattern responses to varying odor concentrations. In the olfactory bulb (OB), odor concentration appears to be encoded through temporal shifts in the spiking latency of mitral and tufted cells, with higher concentrations eliciting earlier and more synchronized responses rather than total spiking output or rate^2,3^. This temporal code is subsequently transformed in the piriform cortex (PC) into ensemble-based representations, where recurrent excitation and feedback mechanisms amplify early inputs from the OB while suppressing later responses, resulting in a multiplexed code for odor concentration^4^. The clear importance of temporal patterns emphasizes the functional role played by neural oscillations in how the olfactory system processes and makes sense of odor information. Studies across species, from amphibians to rodents, have shown that the amplitude and frequency of OB oscillations vary with odor concentration^5–7^. In rodents, increasing odor concentration is associated with enhanced cortical synchrony in the piriform cortex ^8^, while beta oscillations are modulated by odor volatility^9^. These findings suggest that oscillatory activity encodes key aspects of odor concentration and intensity.

Few studies have explored how the human brain process odor intensity. In essence, electrophysiological and imaging studies have revealed that neural responses are more tightly linked to the subjective percept of intensity rather than the mere chemical concentration of odorants^10,11^. Power of obtained event related potentials obtained from the scalp are positively correlated with perceived odor intensity^10^ and perceived odor intensity correlates with activity in the human PC using fMRI^11^. However, no study has examined how the human OB processes odor intensity. Moreover, the electrophysiological response of the PC to variations in odor intensity remains unexplored. As such, the mechanisms by which the early nodes within the human olfactory system communicate intensity-related information is unknown. This knowledge gap is largely due to the lack of non-invasive methods for recording OB activity outside of surgical settings. Recent technological developments have, however, enabled in vivo, non-invasive, and simultaneous electrophysical assessment of the OB and PC functions in humans possible. Specifically, we recently developed and validated the electrobulbogram (EBG); an EEG-based method for assessing oscillatory activity of the human OB and PC^12,13^.

In the present study, we used the electrobulbogram to investigate how the OB and the PC, two central primary nodes, process and communicate odor intensities at its earliest stages. We focus on neural oscillations in response to two key aspects: odor concentration and trial-by-trial perceived odor intensity (Figure 1A). Given that oscillatory burst observed in electrical fields has been proposed as a proxies for population spiking or activity motifs^14^ and that previous animal studies have primarily focused on spiking activity in relation to intensity, we also assessed burst activity in relation to both concentration and perceived intensity to enable more direct comparisons with the existing literature.

**Figure 1.**
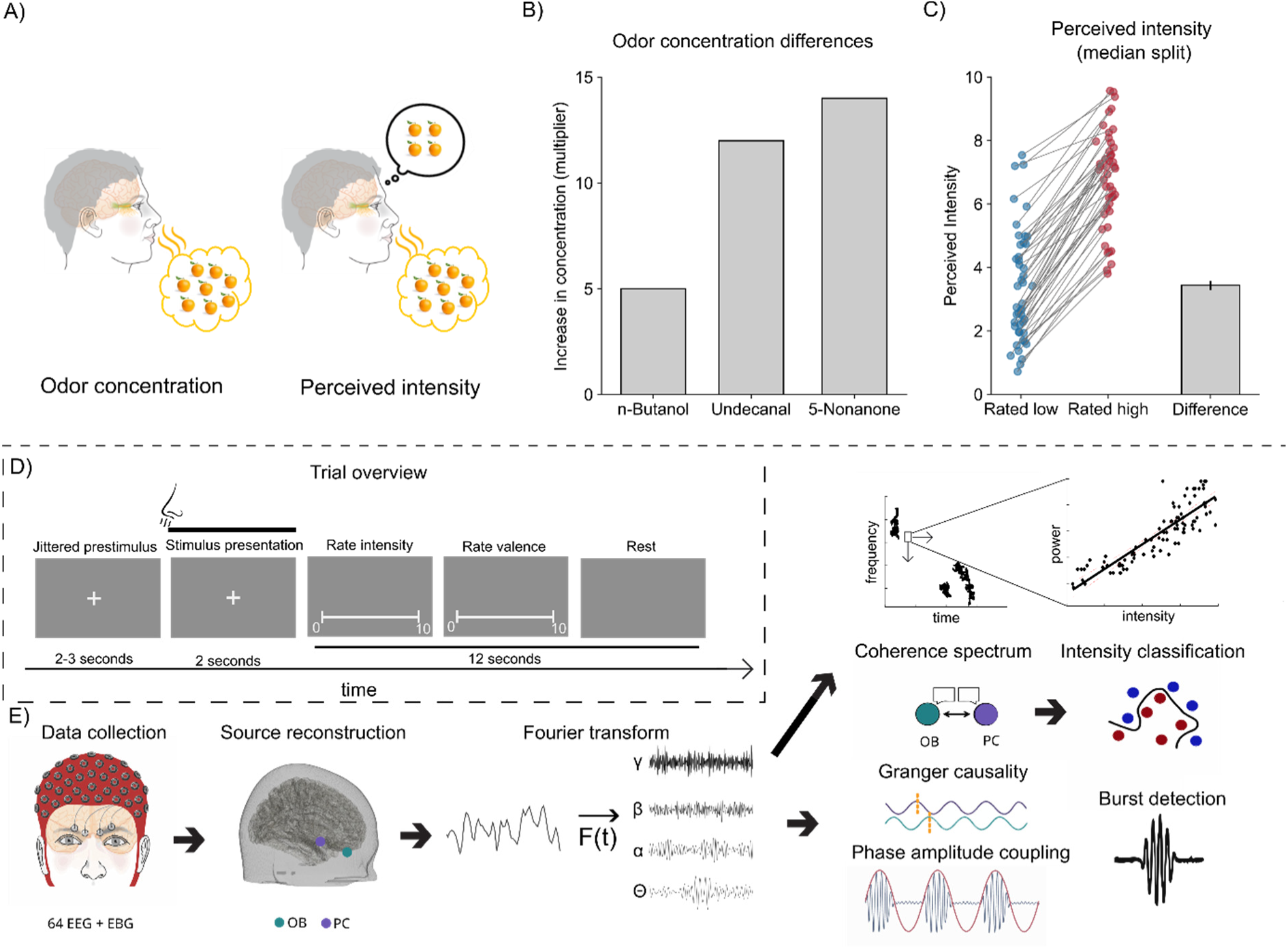
Stimuli characterization, trial overview, and analysis pipeline. **A**) The difference between odor concentration and perceived intensity is that odor concentration is a physical property, measurable through counting molecules, while perceived intensity is what the participant experiences. **B)** Differences between the Low and High concentration conditions for each odor, expressed as multiplier. **C)** Rated perceived intensity based on median split of all individual trials. Each dot represents the mean rating of a participant with their ratings in both conditions connected with line-marking. The bar represents average differences between rated Low and rated High. Blue color denotes Low and red color denotes High and error line in bar represents standard error of the mean (SEM). **D)** Overview of a single trial where the participant is presented first with a jittered pre-stimulus period before the odor is released when the participant breathes in. Odor is then on for 2 seconds and immediately following odor stimulation the participant rates odor intensity and valence on a 0-10 visual analog scale. The rating and the resting time before the start of the next trial is 12 seconds, generating at least a 14-second-long inter-trial interval. **E**) Flowchart outlining the analysis pipeline from data collection to the different methods used to analyze the results.

We hypothesized that the two central primary nodes of the early human olfactory system would be more attuned to the subjective perception (i.e., perceived intensity) of the odor than to the physical concentration of odorants.

## Materials and method

### Participants

A total of 53 individuals participated in the study. Following the exclusion of 7 participants, for reasons outlined in the preprocessing section, the final sample included 46 participants (mean age 30, SD ±8.8; 27 of which were women). Participants with functional anosmia or severe hyposmia were excluded from the study. Olfactory function was assessed using a brief odor identification screening test consisting of five common odors. For each odor, participants were asked to choose the correct label from four options. A minimum of three correct responses was required to meet the inclusion threshold. Participants scoring below this cutoff were excluded to ensure adequate baseline olfactory function. Participants were informed about the purpose and procedure of the study and provided signed informed consent before participation. The study was approved by Swedish National Ethical Permission Board, Etikprövningsnämnden (EPN: 2017/2332-31/1), and performed in accordance with the declaration of Helsinki.

### Testing procedure

Testing was conducted in a sound-attenuated and well-ventilated recording booth, built for odor testing and assuring no-to-limited background odors. During the testing session, participants wore earplugs and headphones that played low-volume white noise to mask any potential auditory cues from the olfactometer, odor delivery setup, or other external sounds. Event timing and stimulus triggering were implemented using PsychoPy 3^15^. Because stimulus presentation was triggered by the individual’s nasal inspiration, participants were instructed to breath normally through their nose throughout the experiment without having to purposely match the odor onset. The experiment consisted of four 15-minutes blocks with a break between each one (Figure 1D).

Each block consisted of 35 trials, resulting in a total of 140 experimental trials (3 odors with Low/High concentration and Clean air, 20 trials each) for each participant. Odors were delivered in a randomized sequence, with their presentation evenly balanced across the testing blocks. Following each trial, participants had to rate with a mouse click the intensity and pleasantness of the stimulus on a visual analogue scale with an underlying scale with 100 scale steps (operationalized as 0.0-10.0). Rating scales were anchored with the text “not perceived/very unpleasant” to “very intense/very pleasant” at opposite ends. The Inter-trial interval (ITI) was jittered but set to at least 14 seconds to limit odor habituation.

### Odor delivery and odor stimuli

Three neutral odors, chosen from different chemical classes, were selected based on extensive in-house testing- were. Each odor was diluted into two different concentrations, resulting in two subsets of odors (Low and High). Using odors from three distinct chemical classes allowed us to draw conclusions that are more likely to generalize across odor processing and to minimize odor-specific effects. In addition, a no-odor condition, i.e. clean air trials in which the ongoing airflow was replaced with a “Clean air” airflow, were presented. All air used in the experiment underwent cleaning stages to remove unrelated odors potentially originating from the compressor system, where the air was passed through both active coal and micro filters. Low-concentration odors were obtained by diluting neat n-Butanol (an alcohol, Fisher Chemicals, CAS 71-36-3), Undecanal (an aldehyde, Sigma-Aldrich, CAS 112- 44-7), and 5-Nonanone (a ketone, Sigma-Aldrich, CAS 502-56-7) to 0.8%, 1.4%, and 1.3% volume/volume in diethyl phthalate (99.5% pure, Sigma-Aldrich, CAS 84-66-2), respectively; while high-concentration odors were obtained by diluting the aforementioned odors by 4% (a 5-fold increase in concentration), 17% (a 12-fold increase), and 18% (a 14-fold increase) in diethyl phthalate, respectively (Figure 1B). Concentrations were selected so that odorants within each subset (Low and High) would be iso-intense; ensured by pilot studies. Odors were delivered birhinally for 2 seconds per trial using a computer-controlled 8-channel olfactometer and presented using a sniff-triggered design^16^. A thermo-pod (sampling rate of 400 Hz; PowerLab 16/35, ADInstruments, Colorado) with a temperature probe inserted just inside participants’ nostrils monitored the breathing pattern based on intranasal temperature. Based on the breathing signal, an individual threshold was set to trigger the olfactometer at the beginning of the inhalation phase to synchronize odor presentation with nasal inspiration to avoid inducing attention-related EEG artifacts. To avoid any potential tactile sensation at the odor onset, a birhinal airflow of 2.7 liters per minute was used for odor delivery and inserted into an ongoing constant airflow of 0.3 liters per minute of clean air. The olfactometer has an onset delay of approximately 150 milliseconds. This delay, representing the time required for the odor to reach the nasal epithelium and dependent on tubing length and airflow, was verified using a photo-ionization detector (Aurora Scientific, Ontario) prior to the beginning of the study. Critically, the established odor delivery delay was taken into account in subsequent analysis, meaning that time point zero in all results corresponds to when the odor enters the nostrils.

Perceived odor intensity is imperfectly related to odor concentration ^1^. Whereas there is a general agreement between a specific odorant’s concentration and how intense it is perceived by a human rater, the percept varies between individual trails based on a plethora of factors, such as intranasal environment, perceived intensity of prior trails, adaptation and habituation, sniff behavior, sampling duration, rating behavior, trigeminal components, attention, to mention a few. This means that over the course of an experiment, a given concentration of an odor can demonstrate a range of perceived intensities that is a true percept and not solely attributable to rating behavior. Seen over all trails, the distribution of rated perceived intensity for the six odorant conditions we used produced intensity percepts spanning nearly the full rating range with a slight overrepresentation around predicted Low/High concentrations (Supplementary Figure 1). To assess neural responses to perceived odor intensity, as outlined below, we use trial-by-trial ratings. Average min-max range of ratings across participants was 8.28 ± 1.72 (Supplementary Figure 2), demonstrating a good range of ratings. For our support vector machine classification analyses, which require a binary factor, we used for each individual a median split to create low and high rated intensity trial, independent of stimuli. Here, the low rated intensity trials were on average rated as 3.36 (± 2.14 SD) and the high rated intensity trials were on average rated as 6.80 (± 1.86 SD), *t*(45) = 23.33, *p* < 8.93e-27, an average increase between Low and High in perceived intensity with 102% (Figure 1C).

### EEG and EBG recordings

Continuous neural activity was recorded at a frequency of 512 Hz using 64 EEG and 4 EBG active electrodes (ActiveTwo, BioSemi, Amsterdam, The Netherlands), Figure 1E. EEG electrodes were placed following the international 10/20 system and EBG electrodes were placed on the forehead slightly above both eyebrows. Position of each electrode was digitalized in stereotactic space using a neuronavigation system (Brain-Sight, Rogue Research, Montreal, Canada). The digitalization protocol involved localizing fiducial landmarks (nasion, left and right preauricular points) and the central point of each EEG/EBG electrode. The identified landmarks were used to map each electrode to the standard MNI space. In subsequent analysis, the recorded electrode coordinates were used in the eLORETA algorithm to determine the sources of neural signals, described below.

During recording, the signal was high-pass filtered at 0.10 Hz and low-pass filtered at 100 Hz within the ActiView software (BioSemi, Amsterdam, The Netherlands). Prior to the actual EEG/EBG recording, the impedance of each electrode was visually inspected and electrodes with an offset greater than 40 mV were manually adjusted until meeting a value below the accepted threshold.

### Signal processing and analysis

#### Preprocessing

Prior to preprocessing, five subjects were removed due to a too small variability in the rated perceived intensity in that they rated all odors as basically the same. For the remaining 48 participants, data was epoched to 5000 ms long segments, from 1000 ms pre-stimulus to 4000 ms post-stimulus, and re-referenced to the average activity of all electrodes. Data were then high pass filtered at 1 Hz and a notch filter at 50 Hz was applied to reduce power line interference. Faulty electrodes were detected through visual inspection and corrected using spline interpolation. Trials with large muscle movements were detected and removed by extracting z-scored Hilbert transform amplitude values with a threshold of 7. Furthermore, eye blinks were removed with Independent Component Analysis with the InfoMax algorithm. Two participants with less than half of the trials left after this step were removed from further analysis. After preprocessing, the remaining and final 46 participants had an average of 123 ± 6.84 trials left for final analyses.

### Source time-course reconstruction

Source time-course for the OB and PC dipoles was reconstructed through a conduction model based on structural MRI of the participant’s head. The structural MRI was segmented into five materials CSF, gray matter, white matter, scalp, and skull with conductance’s [0.43, 0.01, 1.79, 0.33, 0.14]^17^. The digitized electrode positions were co-registered to the headmodel and the forward problem was solved using the Finite Element Method through simbio^18^.

A cortex model built on icosahedrons with resolution 7 generated in Freesurfer based on individual structural MRI was used as a source model. This source model was then transformed into MNI stereotactic space to ensure that identical regions are used for all participants. This source model was used to attain the source activity through solving the inverse problem with eLORETA with a regularization parameter set to 10 % with Singular Value Decomposition used to project the source activity along the principal axis. The analysis was later constrained to four ROIs where the dipoles correspond to left *(x−4, y+40, z−30)* and right OB *(x+4,y+40,z−30)*, determined based on T2 weighted images, as well as left *(x -22, y+0, z - 14)* and right PC *(x+22, y+2, z -12)*^19^. The source reconstruction was performed with Fieldtrip toolbox 2023 within Matlab 2023a^20^.

### Linear modelling of the intensity percept

To evaluate how odor concentration and intensity is processed within the OB and PC, frequency spectrums were generated using the Superlet method^21^ with width 8 and an adaptive order based on frequency bands. This approach was taken to achieve a high resolution in both time and frequency. To assess the relationship between power amplitude and odor concentration as well as perceived intensity, a Mixed Effect Model was applied to each time-frequency point in the spectrum. The spectral power was used as the outcome variable while perceived intensity or concentration, together with perceived valence, amplitude of the breathing signal and area under the curve (AUC) of the breathing signal, were used as fixed effects. We included perceived valence and breathing parameters to control that potential neural effects would not be linearly related to either breathing or perceived valence. A random intercept was defined on the subject level according to the formula (power ∼ intensity + valence + breathing amplitude + breathing AUC + (1|subject)) or (power ∼ concentration + valence + breathing amplitude + breathing AUC + (1|subject)).

### Source connectivity

Both functional and effective connectivity approaches can be taken when investigating the connectivity between neuronal populations. While amplitude and phase relationships are evaluated through functional connectivity^22,23^, effective connectivity instead determines the predictive relationship between the two^24^. To get an understanding of when two neuronal populations are connected, and which one of them causes the other, both approaches are preferred.

The functional connectivity was evaluated through the coherence spectrum, allowing us to evaluate whether there is a linear transfer of information between the OB and PC. OB-PC coherence spectrum was calculated through Fourier transforming the source reconstructed time-course with the previously mentioned Superlet method. Effective connectivity was evaluated through spectral Granger causality. Granger causality assesses whether the future of a time series (X) can be predicted by past values of X alone or if it is more accurately predicted by past values in the alternative time series (Y)^25^. Spectral Granger causality builds upon the same concept, but the assessment is here performed in Fourier space. To this end, we applied a multi-tapered fast Fourier transform with a DPSS taper based frequency smoothing of 3 Hz. The Fourier transform was applied to the whole stimulus period and averaged over both hemispheres to increase statistical power. Statistical significance was determined on a group level by applying a two-tailed Student’s *t*-test.

### Support Vector Machine classification

A Support Vector Machine (SVM) classifier was applied to the trial averaged OB-PC coherence spectrogram to determine whether any significant transfer of information related to either perceived intensity or odorant concentration could be decoded. The entire coherence spectrogram was analyzed in a binned searchlight manner where the bins were 12 Hz and 100 ms. Bins with less than 10 neighbors were excluded from further analysis. A leave-one-out scheme was applied on the data where each participant was left out once per squared area. The accuracy on the group level was compared with a distribution of 1000 classification results where the labels were shuffled. A Monte-Carlo simulated one-sided non-parametric *t*-test was used to evaluate significance and significant clusters were evaluated with the WCM algorithm^26^.

### Phase-amplitude coupling

The integration of neural activity in the brain is believed to be facilitated by a phenomenon wherein the phase of lower-frequency electrophysiological oscillations modulates the amplitude of higher-frequency oscillations, so-called phase amplitude coupling (PAC)^27^. Our past studies have demonstrated that perceptual information is communicated back to the OB from the PC in the beta band and subsequently processed within the gamma band in the OB^28^. Therefore, to evaluate information transfer in between frequency bands, we assessed PAC based on the tort algorithm^29^ between the beta and gamma band in the timepoints where our analyses in the coherence spectrum demonstrated information transfer from the PC to OB. A time window of 150 ms was used for the PAC-analysis based on the duration of the beta activity in the coherence spectrum. The trial level modulation index (MI) derived from the PAC was then included as the response variable in a linear mixed-effect model with perceived intensity, perceived valence, breathing amplitude, and AUC of the breathing signal as fixed effects and a random intercept based on participant. The significant clusters were evaluated with a 1000 permutations Monte-Carlo test with WCM cluster correction.

### Burst analysis

To investigate oscillatory bursts in response to odor stimulation, we analyzed time-frequency representations of the neural activity recorded during odor perception where bursts were identified as oscillatory activity in the power spectrum exceeding 1 standard deviations above 10-trials mean and lasting for at least three cycles of the band’s average frequency^30^. Because of their intermittent nature, burst events were determined at the single trial level and extracted for a window of 2000 ms following stimuli onset. For each participant, we first calculated the burst rate separately for OB and PC and the two frequency bands of interest (beta [12-30 Hz], and gamma [30-100 Hz]). Burst rates were calculated across trials and in some analysis combined across participants returning a grand average burst rate, per frequency band and region. For the main analysis, we kept single trial data in order to examine the relationship between burst activity and behavior parameters (perceived intensity, valence, and odor concentration). We subsequently applied linear mixed-effects models using separate models for beta and gamma bursts in OB and PC. Participants were included as random effects to account for inter-individual variability, and t-statistics for the fixed-effect predictors were used to determine their significance in influencing burst dynamics.

## Results

### Gamma and beta amplitude relates linearly to perceived odor intensity in the OB and PC

We first determined whether odor concentration or perceived odor intensity is related to gamma and beta amplitude by assessing each time-frequency point in the power spectrum with a linear mixed-effect model. To control for individual differences in rating behavior, we included Participant as an intercept, and to control for the influence of perceived odor valence and breathing, we included perceived valence as well as the breathing parameters amplitude and AUC as fixed effects in all models. Models were statistically assessed with a 1000 permutations Monte-Carlo test using WCM cluster correction.

When we assessed odor concentration, we found only minor areas within the time-frequency spectrum where concentration related to power amplitude (Figure 2A-B). None of these areas survived cluster correction. However, when assessing perceived odor intensity, there were cluster corrected areas in both the OB and PC beta bands where rated intensity was found to be related to amplitude. In the OB, we found that intensity ratings were related to power in the beta band (12-25 Hz) in a time span from 600 – 1600 ms *t = 3.51*, *p* = 0.001, CI = [.0081,.0101] (Figure 2C). In the PC, a similar area of activity was found somewhat later (850-2000 ms) in the beta band between 15-30 Hz, *t* = 3.03, *p* = .004, CI = [.0142, .0222] (Figure 2D).

**Figure 2.**
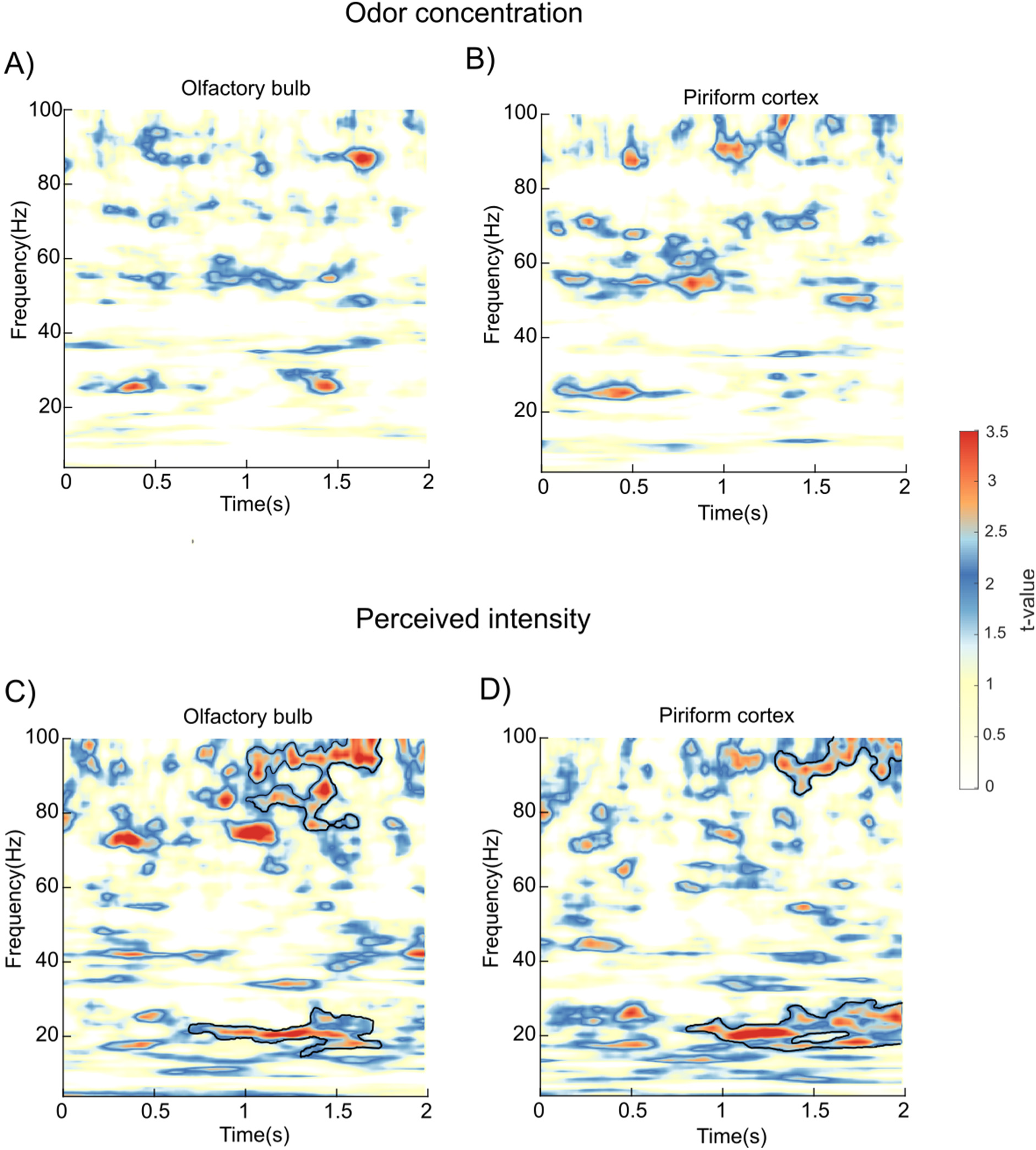
Gamma and beta power are predicted by perceived intensity, but not actual odor concentration, in OB and PC. **A)** Mixed effect model applied to OB spectrum where concentration (High or Low) predicts power amplitude. Valence and breathing parameters are included in the model and hence controlled for, and the intercept is adapted based on the subject. Significant areas after cluster correction are marked with a black border. **B)** Same as A, but for PC. **C)** Same model as in A but with perceived odor intensity instead of odor concentration. **D)** Same as A but for perceived intensity in the PC.

While the area included in the significant cluster starts earlier in the OB, there were intermittent significant peak activity both in the OB and PC beta band already 200 ms after odor onset, albeit not surviving statistical cluster corrections. Furthermore, the strength of the association is higher in the PC beta areas around one second into the trial, *t* = 4.65, *p* = 3.36e-6 , CI = [.0757, .186]), compared to the OB, *t* = 4.08 , *p* =4.49e-5 , CI = [.0726, .210], suggesting that at this time point, the PC is more informed about the perceived odor intensity compared to the OB.

When we assessed the gamma band, similar results emerged. Clusters in the OB gamma band amplitude were related to perceived odor intensity around 1100-1800 ms after odor onset, which spanned a wide frequency range (70-100 Hz). Similarly, clusters in the PC gamma band emerged at a slightly later timepoint, 1400-2000 ms after odor onset, in the 80- 100 Hz frequency range. The relationship between perceived intensity and power amplitude were similarly strong in the PC compared to the OB, *t* = 2.37, *p* = .022, CI = [.020, .064]; and *t* = 2.41 *p* = .02, CI = [.021, .061], respectively. Similar to the beta band, we found significant areas at early timepoints of the spectrogram, especially for the OB. Already 200 ms into the trial, we obtain significant links between power and perceived intensity around 70-80 Hz. This result does not survive the cluster correction; however, it aligns with our *a priori* hypothesis that gamma band hold information related to odor intensity in the OB at early time points. If we predict that we will find gamma activity between 100-500 ms into the trial and evaluate the area in this time frame, result do survive cluster correction.

Past work in animal models suggests that the OB process odor concentration^5–7^. However, there was a surprising lack of clear effects in our analysis of the time-frequency spectrum in the OB. To assess whether there are latent parameters of OB processing of odor concentration that our method is not able to extract, we performed principal component analyses of the full raw OB time series, separately for the high and low concentration condition, and plotted over the full odor presentation time-period (0-2000ms). There was a clear separation between OB processing of high vs low concentration early on in the trajectory, only to subsequently coalesce (Supplementary Figure S3). This suggests that there is concentration processing-dependent processing of odor concentration that is not present in the time-frequency domain.

We obtained a broad range of intensity percepts for nearly all participants. To assess whether the obtained range is linked to perception and not primarily due to variation in rating behavior, we utilized the established fact that there is a weak but linear relationship between perceived intensity and the sniff response. We found that subjective odor intensity ratings positively correlated with the area under the curve (AUC) of the inhaled odor, *t =* 3.79, *p* = .00015, while odor concentration (High/Low) did not, *t* = .19, *p* = .85. Note that in all of our other analyses, sniff parameters are included as parameters of no interest.

In our analysis, we used a dichotomized parameter for analyses related to odor concentration (High/Low). It can be argued that the lack of allowed variance masks potential results in the mixed effects models we deployed. To assess this, we transformed our six odor conditions into a measure of effective concentration based on their known vapor pressure multiplied by volume concentration (dilution), producing a linear scale of theoretical effective concentration with 6 scale steps. We then perform identical analyses as described above for the first analyses, but replacing the dichotomized parameter with this new linear parameter. The obtained result for effective concentration (Supplementary Figure S4A-B) was similar to what we obtained for concentration (Figure 2A-B), i.e., no larger significant area evident in the obtained time-frequency spectra.

### Perceived intensity is reciprocally communicated between the OB and PC in the gamma and beta band

We then wanted to determine how odor concentration and perceived odor intensity is communicated between the OB and PC. To this end, we first calculated the coherence spectrum between the two nodes and subsequently applied two separate SVM classifiers for binary classification with either high and low concentration or high and low perceived intensity, based on a median split of the ratings. There was no area in the time-frequency matrix where our classifier managed to extract enough information from the oscillatory activity to correctly classify odor concentration (Figure 3A). In contrast, our classifier could extract transmission information between the two nodes that allowed it to correctly classify perceived odor intensity at multiple locations (Figure 3B). First at an early stage, 200 ms after odor was delivered at the nose, in the higher gamma band (70-90 Hz), *t* = 2.56, *p* = .014, CI = [.0068, .0212]. The confusion matrix, Figure 3E-I, demonstrates that in this specific region, the classifier was better at classifying low perceived intensity than high, achieving 76% accuracy compared to 67%. This early gamma region furthermore corresponds to activity that was found when applying linear mixed-effects models on the power spectrum of the OB and PC (cf. Figure 2C-D). These regions in OB and PC did not survive cluster correction; however, the areas were significant on voxel level and the alignment of findings indicate that both OB and PC process perceived odor intensity already at this stage.

**Figure 3.**
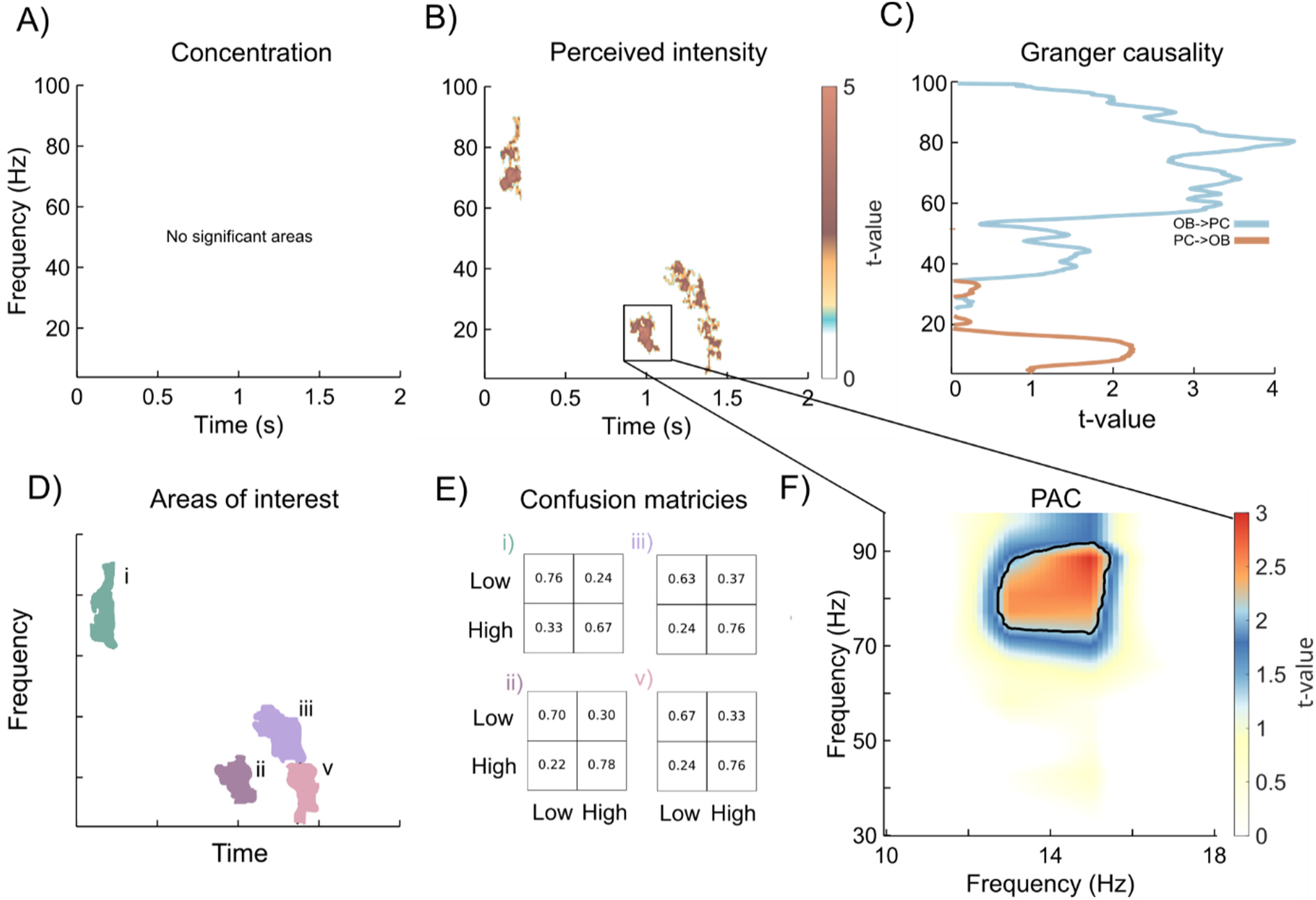
Perceived odor intensity is communicated between the OB and PC. **A)** No significant activity was found when classification of high and low concentrations was applied to the coherence spectrogram. **B)** Classification of high and low intensity in the coherence spectrogram demonstrates significant classification accuracy in the gamma and beta band. **C)** Granger causality show communication from OB to PC in the gamma band and from PC to OB in the beta band. Indicating that the beta region in B is in the direction from the PC to OB. **D)** Areas of interest from B. **E)** Confusion matrices describing the four significant areas in B. **F)** PAC for the time window 950-1100 ms show that the beta band modulates the gamma band in the OB at the time where perceived intensity is communicated back from the PC to the OB.

The later beta activity (12-25 Hz), around 1 second into the trial, *t* = 2.56, *p* = .014, CI = [.0068, .0212], also aligns with activity found in the linear mixed effects models of the OB and PC where both have clusters in this region that significantly predicts OB and PC oscillatory power based on perceived intensity. This is also the region with the highest classification accuracy (74%), with better performance in classifying high perceived intensity (78%) compared to low (70%).

The last significant region in the resulting time-frequency matrix spans between 1200-1500 ms and stretches from beta to lower gamma (10-38 Hz, *t =* 3.03 *p* = .004, CI = [.0001, .0079]. We therefore decided to divide the cluster into beta and gamma bands when looking at the confusion matrices. In both bands, the classifier demonstrated a stronger ability to classify high perceived intensity (76% in both beta and gamma) compared to low perceived intensity, classified at 67% in beta and 63% in gamma.

Odors were selected to be neutral in valence and therefore displayed little variance (Supplementary Figure S5A). Moreover, perceived valence was included in analyses as a factor of no interest. That said, to assess whether our classification in the coherence spectrum might have been mediated by perceived odor valence, we conducted a classification analysis for perceived valence (high/low rated valence using a split-half on trial and individual level procedure) in the coherence spectrum. As can be seen in Supplementary Figure S5B, results showed no overlap between the regions identified for valence and those associated with perceived intensity.

We then assessed the direction of information transfer by conducting a frequency-resolved Granger causality analysis where we contrasted the direction from the OB to the PC with the direction from the PC to the OB to remove joint variance. We found that information from the OB to PC appears to be reflected in high gamma band activity, *t* = 5.04, *p* = 8.3e-6, CI = [0.2275, 0.5313] (Figure 3C). In contrast, the lower frequency components appear to reflect communication from the PC back to the OB, *t* = 2.46, *p* = .0134, CI = [0.0833, 0.8417]. This suggests that the beta-band activity observed around one second in the coherence spectrum may be linked to information transfer from the PC to the OB. Therefore, based on the hypothesis that the PC convey updated information of the percept to the OB and that the OB would update its internal state based on this information, we performed a PAC-analysis with the time and frequency window of this area as the region of interest and found modulation of the beta (12-16 Hz) phase to the gamma (70-90 Hz) amplitude related to perceived odor intensity, *t* = 2.35, *p* = .023, CI = [.0137, .0323] (Figure 3F). Moreover, Granger causality analysis suggests that the beta region found in the coherence spectrogram of perceived intensity around 1400 ms is within the area of significant information transfer from the PC to the OB and would therefore indicate continuous information transfer between these regions.

### PC beta and OB gamma bursts indicate information transfer

Analyzing intermittent burst activity provides additional insights beyond those offered by traditional wave-based analyses because it considers that oscillatory activity is intermittent and varies from trial to trial. This approach aligns with the view that cognition and executive control are supported by rapid transitions between discrete, transient neural states, rather than by slow, continuous dynamics^31^. Furthermore, oscillatory bursts are closely linked to spiking activity, making them a useful proxy for population-level spiking or activity motifs. Thus, to further investigate the relationship between odor intensity processing, cognition, and neural oscillations, we analyzed single-trial burst activity in the beta and gamma band during odor presentation, examining its association with both odor concentration and perceived intensity. We extracted burst activity in each broader frequency band and assessed the relationship between burst counts and perceived intensity and odor concentration using a linear mixed-effect model. The response variable was transformed with a Box-Cox transformation to fulfill the requirements of normality, and model outcome was statistically evaluated with a 1000 permutations Monte-Carlo test with WCM cluster correction.

When relating odor concentration to gamma and beta activity in the OB, we found no significant relationship between OB beta burst behavior that withstand cluster correction. We did find, however, at an uncorrected level (*p* < .05) that beta burst rate related to odor concentration from 1500-1800 ms into the trial *t* = 2.80, *p* = .0051, CI = [7.70e-5, 5.85e-4]. A similar pattern was observed in the PC at the same time, albeit also at uncorrected significance levels, *t* = 2.87, *p* = .004, CI = [1.50e-4, 7.94e-4].

When we instead related gamma and beta burst behavior to perceived intensity, we found significant cluster corrected burst activity in the gamma band within the OB at two time points. First, around 800-1100 ms after odor onset, *t* = 2.88, *p* = .006, CI = [.0012,.0108]. Interestingly, after this event, the OB gamma burst activity then became anti-correlated to perceived odor intensity, suggesting suppression of high frequency bursts by intensity. Later, burst activity in the gamma band was again positively related to odor intensity ratings between the timepoints 1350-1450 ms, *t* = 2.53, *p* = .015, CI = [.0075, .0225].

In the PC, significant cluster corrected beta burst activity was found in relation to perceived intensity at a time corresponding to the initial gamma band burst activity in the OB (800- 1100ms after odor onset), *t* = 2.28, *p* = .027, CI = [.0170, .0370]. Critically, this corresponds to the time window reported above when the PC communicates perceived odor intensity back to the OB in the beta band, and where the OB exhibits a phase amplitude coupling between the beta and gamma band connected to perceived odor intensity.

## Discussion

We here demonstrate that oscillatory activity within the OB and PC is robustly associated with perceived odor intensity, but only weakly with the physical concentration of the odorant. Approximately, 200 ms after odor onset, the OB initially broadcasts odor intensity information via fast gamma-band oscillations (∼70–90 Hz) to the PC in a bottom-up manner. Subsequently, around 1 second post-inhalation, the PC returns a top-down signal to the OB in the beta-band (∼15–30 Hz), conveying a more refined representation of the odor’s perceived intensity. This beta feedback modulates ongoing OB activity through phase–amplitude coupling, likely updating the OB’s gamma activity with the enriched perceptual information. Notably, these dynamic interactions were absent when considering physical odor concentration alone; oscillatory power and OB–PC coupling were strongly linked to how intense an odor was perceived, not the stimulus concentration. This reveals that subjective perceptual intensity, rather than raw stimulus properties, dominates neural processing in early human olfactory circuits. Moreover, this suggests that the early olfactory system in humans is tracking the brain’s internal estimate of stimulus intensity; an estimate that may incorporate not just concentration, but also factors like sniff volume, odor sorption dynamics, and short-term adaptation, consistent with prior findings e.g^e.g., 3^. However, even when using an "effective concentration" metric (i.e., incorporating vapor pressure and dilution to linearize physical concentration across odorants), we showed that neural dynamics aligned with these subjective ratings. This does not, however, imply that the OB is oblivious to concentration because our model-free raw signal analyses indicated early processing differences between concentrations, but this did not manifest itself in any oscillatory activity.

We found a strong link between OB processing and subjective perception, suggesting that by the time oscillatory population activity is measured (hundreds of milliseconds post-stimulus), it reflects a perceptually transformed representation. This view is reinforced by rodent studies showing that the OB can encode an internal variable corresponding to perceived intensity rather than just linearly reporting concentration^3^, and that intensity coding can involve specific population dynamics^32^. The striking dissociation between the encoding of perceived intensity and physical odor concentration in OB and PC oscillatory signatures underscores the brain’s active role in transforming raw sensory input into a behaviorally relevant perceptual scale.

We here offer insight into how the human olfactory system might implement intensity coding, which in animal models involves complex temporal dynamics rather than simple rate codes. For instance, in the mouse PC, overall neural firing rates are relatively concentration-invariant for a given odor identity, but intensity information can be extracted from temporal features of the population response^2^. Specifically, piriform cortex uses a dual-component code where an initial subset of neurons responds rapidly and largely independent of concentration (carrying odor identity), while a later subset responds with latency shifts as concentration changes, thus conveying odor intensity^2^. Notably, this dissociation means that odor intensity is not simply represented by “more firing” at higher concentrations, but rather by dynamic timing patterns; a principle also found in insects and fish^7,33^. Our findings provide a potential mechanistic basis for this type of coding in humans. The early, bottom-up OB-to-PC gamma signal (∼200 ms) might relate to an initial, somewhat concentration-invariant odor representation, while the later PC-to-OB beta feedback (∼1 s), which we found to be strongly tied to the perceived intensity, could reflect the processing of this intensity-varying component. Consistent with this, PC beta-band activity showed a stronger correlation with the magnitude of perceived intensity compared to OB activity at this later time point, suggesting cortical integration of sensory input and contextual/cognitive factors to produce a refined intensity code. This aligns with the piriform cortex’s capacity, via recurrent circuitry, to amplify or transform OB input^2^ and with proposals that olfactory beta oscillations signify network-level coupling involving cortical feedback^28,32,34^. Indeed, beta oscillations have been hypothesized to bind distributed olfactory areas into coherent network states^35^, a function precisely aligned with a top-down PC-to-OB signal refining the intensity percept.

Our observation that gamma oscillations mediate bottom-up transfer while beta oscillations mediate top-down feedback resonates strongly with theoretical and experimental work in animal models. It has long been proposed that fast gamma oscillations primarily reflect local circuit processing within the OB and the initial feedforward relay of olfactory information, whereas slower beta oscillations engage a larger loop including the olfactory cortex and potentially higher-order areas^34,36^. For example, OB beta oscillations are typically weak or absent under anesthesia or when cortical feedback is disrupted, while gamma oscillations persist, indicating that beta rhythms critically depend on an intact OB-PC loop^36,37^. Our human data, demonstrates significant Granger-causal influences from OB to PC in the high gamma-band and conversely from PC to OB in the beta-band during intensity processing, support this model. Critically, these findings align and complement recent human research demonstrating decoding of both odor identity (35–45 Hz at around 100 ms) and perceived odor valence (55– 65 Hz at around 150 ms) in the OB-PIR connectivity, both at earlier time points and different frequencies compared to intensity perception^13,28^.

The sequential processing of odor intensity mirrors a two-stage scheme proposed in olfaction and other sensory systems^13,28,33,38,39^, wherein an initial feedforward sweep is followed by feedback that fine-tunes the representation, fitting with the timing of odor-evoked gamma and beta oscillations found in human intracranial PC recordings^40^. This OB-PC loop could potentially be viewed through a predictive coding lens cf^cf. 41^, where the cortex sends predictions to early sensory areas, and discrepancies (prediction errors) drive updates of internal representations to refine perception^42,43^. The observed gamma/beta cycle could instantiate such a mechanism, iteratively minimizing the mismatch between expected and actual intensity.

Evidence that the internal representation of the OB is updated is found in single-trial analysis of our data showing intermittent oscillations^31,44,45^, and that the rate and timing of bursts in the beta and gamma bands were systematically related to perceived intensity. Specifically, OB gamma-band bursts showed a biphasic positive relationship with higher perceived intensity (an initial surge at ∼0.8–1.1 s, followed by suppression, then a later surge at ∼1.35–1.45 s). Crucially, PC beta-band bursts occurred in nearly the same early time window (∼0.8–1.1 s) as the OB’s initial gamma burst surge. This temporal coincidence, coupled with our finding that beta phase in the PC modulated OB gamma amplitude (PAC) during this period, suggests a sequential interaction: as the PC evaluates intensity (reflected in a PC beta burst), its feedback triggers or "gates" OB gamma bursts, thereby updating the OB’s representation in discrete epochs. This aligns with emerging views of cortical function where information is processed in transient "packets" or bursts, facilitating efficient communication and plasticity. Gamma bursts are thought to reflect synchronized spiking in local networks, while beta bursts often involve more distributed networks^34,45^. Thus, a PC beta burst, possibly influenced by higher-order inputs, could prime piriform ensembles to send temporally precise volleys of activity that manifest as synchronized gamma bursts in the OB, ensuring robust and adaptive population coding of intensity. Such within-sniff updates, occurring over hundreds of milliseconds, are sufficiently fast to influence perception on an ongoing basis.

In summary, the human olfactory bulb and piriform cortex engage in a dynamic, frequency-specific dialogue to represent perceived odor intensity. Here even the most fundamental aspect of perception becomes an active, dynamic, and contextually informed process. Initial bottom-up gamma oscillations convey early sensory information, which is subsequently refined by top-down beta-band feedback from the cortex, implemented through PAC and coordinated neural bursts. This framework highlights the primacy of subjective perception in shaping early olfactory processing and suggests that, similar to other sensory systems, olfaction relies on iterative network interactions to construct a stable and behaviorally relevant representation of the world.

## Data availability

All anonymized data and scripts to reproduce the results are available at https://osf.io/t7hj2/

