## Supplementary for "Olfactory bulb and cortex activity reflects subjective odor intensity perception rather than concentration"

---

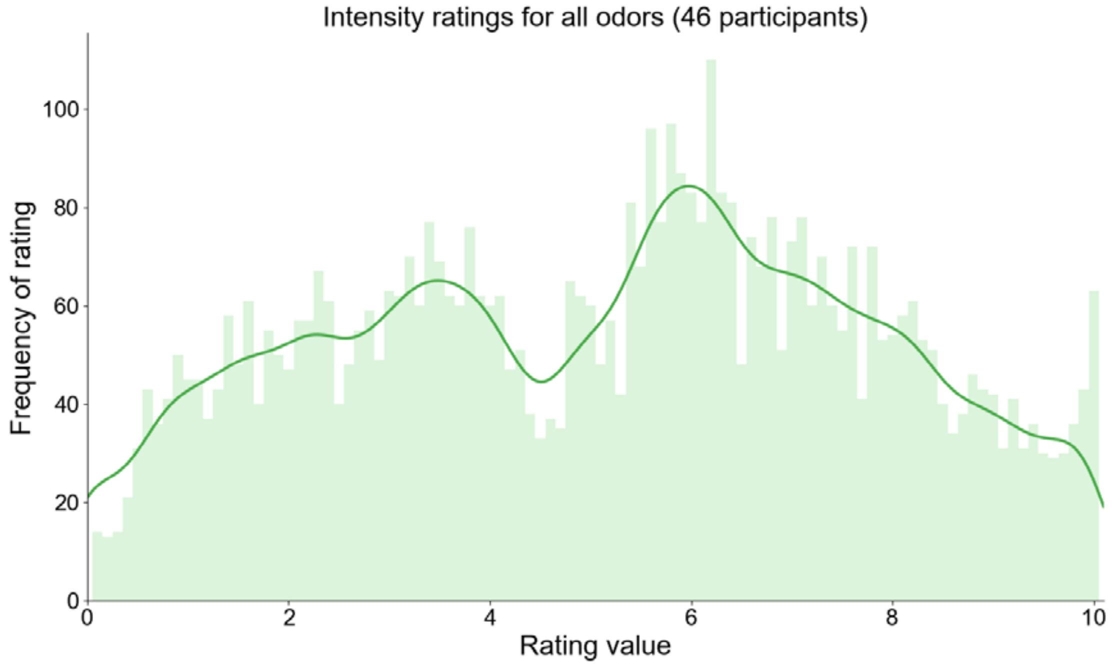


**Supplementary Figure S1:** Histogram of perceived odor intensity ratings for all trials in all participants. Solid line represents the density curve (kernel bandwidth=0.1). This demonstrates that ratings were evenly distributed across the scale with an overrepresentation of ratings at two scale points, in-line with the two odor concentrations used.

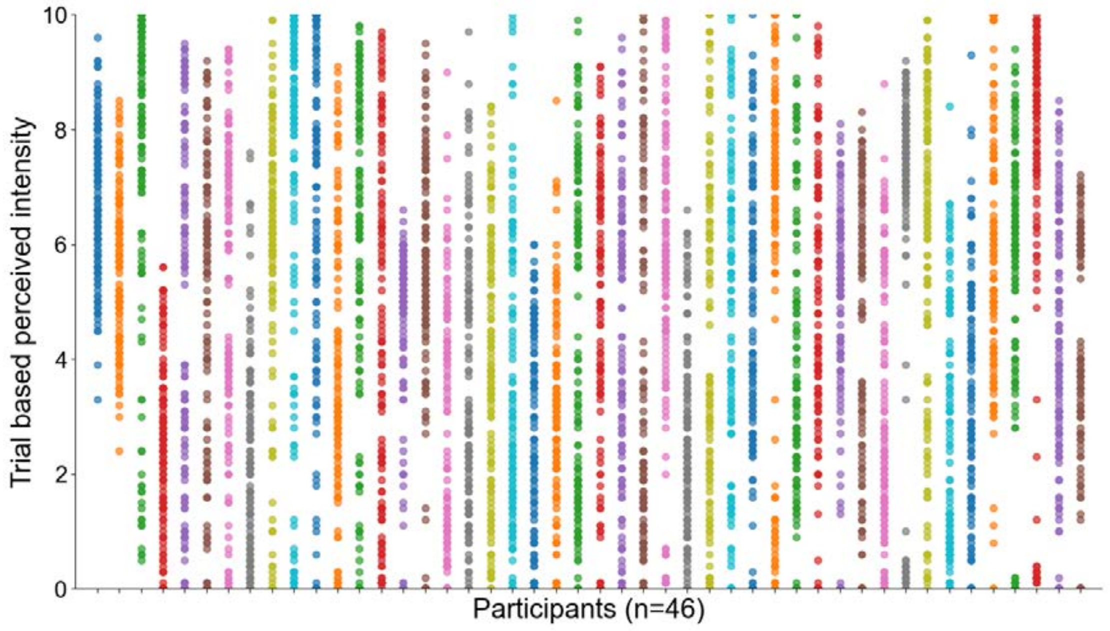


**Supplementary Figure S2:** Odor intensity ratings for each individual trial, sorted diagonally within each participant according to participant number (assigned in the order of entering into the study). This demonstrates that participants experienced odor intensities across the scale and that there is no systematic skewness of ratings across study time. Colors in figure have no meaning other than to visually differentiate individual participants.

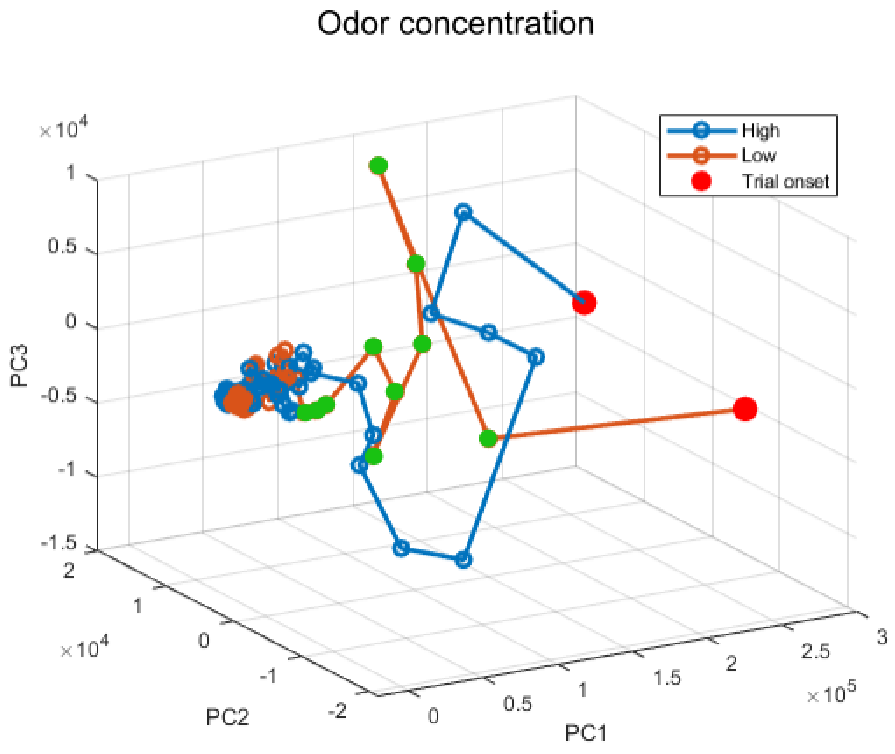


**Supplementary Figure S3:** Trajectory of the power spectrum in the first three principal components, extracted from the signal source data from the olfactory bulb. These results suggest that there is variance in the odor concentration data that we do not capture with our linear statistical models. Green dots represent time-points where we could distinguish between high and low concentrations.

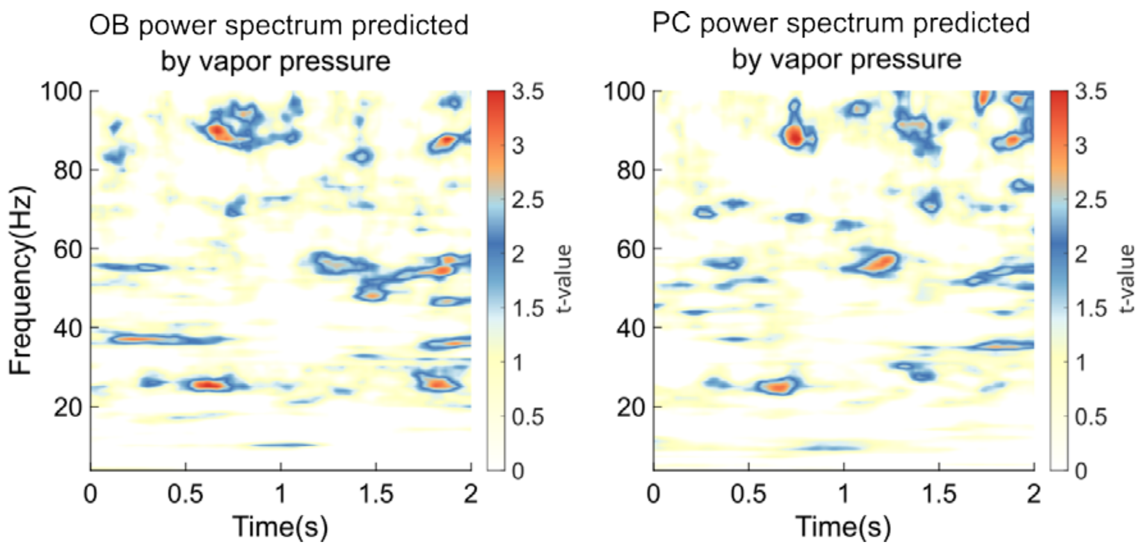


**Supplementary Figure S4:** Similar to Figure 5, but with effective concentration, here denoted as the dilution multiplied by the odors vapor pressure. There was no significant activity for odor concentration across the time/frequency spectra. This obtained result is similar to what was obtained using the dichotomous division into high and low.

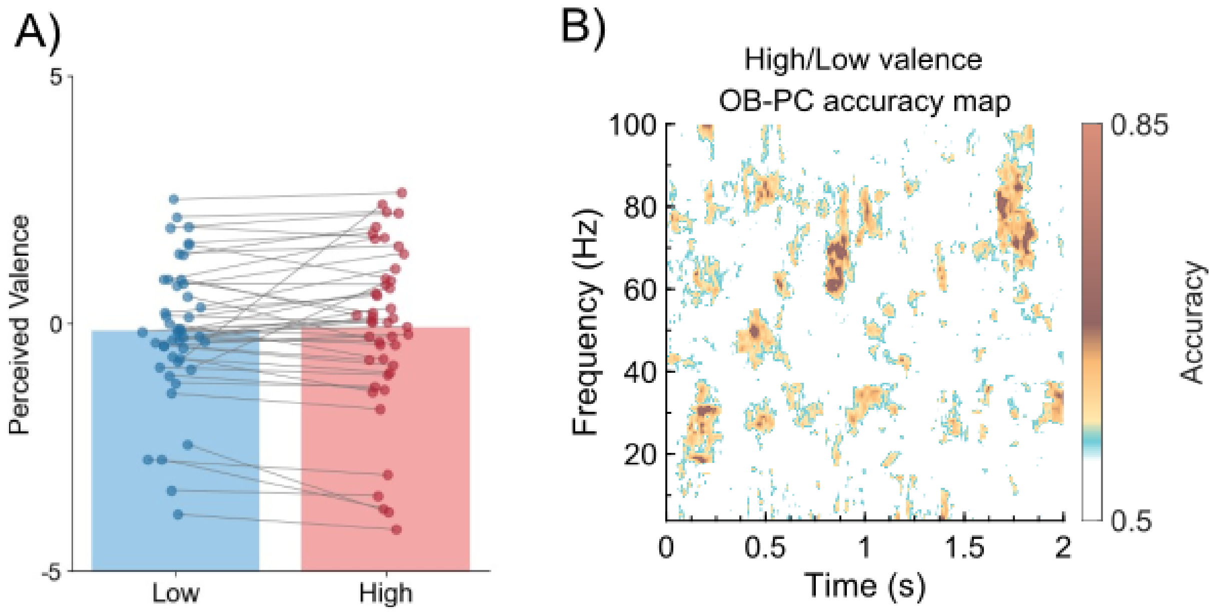


**Supplementary Figure S5: Classification of pleasant and unpleasant as a control**. **A)** Perceived valence did not vary between the two concentrations. High numbers denote pleasant, and low numbers denote unpleasant, perceptual ratings. **B)** None of the areas found overlaps with the areas found in the classification of Intensity.
